# *In vitro* biofilms of *Clostridioides difficile* undomesticated strains: Morphology and properties according to strain diversity

**DOI:** 10.1101/2024.10.18.618822

**Authors:** Anaïs Lemaire, Ludovic Bridoux, Vlad Costache, Aline Crouzols, Frédéric Barbut, Sandrine Petry, Isabelle Poquet

## Abstract

*Clostridioides difficile* is a Gram-positive, spore-forming, obligate anaerobe and an entero-pathogen representing a One Health problem. *C. difficile* infections are difficult to treat and recurrences frequent. *C. difficile* biofilms may play a role in persistence. We studied the biofilms formed *in vitro* by a collection of *C. difficile* undomesticated strains (n=27, 20 toxigenic and 7 non-toxigenic) of 11 PCR-ribotypes (WRT) and isolated from equids. After growth in rich BHIS medium supplemented with glucose for 48h, the biomass of biofilms stained by crystal violet, i. e. adhesive to the surface, was measured, while intact biofilms were observed *in situ*, by confocal laser scanning microscopy, to determine their total biovolume and 3D morphology. All biofilms were relatively similar, even though adhesive, stained biofilms of non-toxigenic strains of ribotype 009 significantly displayed highest biomass. These results suggested that strains of ribotype 009 were able to form a highly cohesive and polystyrene-adhesive biofilm than any other strains. 8 common, representative ribotypes in equids, including 6 clinically important ribotypes in humans, were then selected and one strain chosen for each of them. Their biofilms and planktonic cultures grown under the same conditions were compared for sporulation, toxin production and susceptibility to vancomycin at a highly inhibitory concentration (8 MIC). In both the biofilm and planktonic culture of all selected strains, sporulation was low and toxin production undetectable. The addition of vancomycin at T0=24h followed by a 24 hours exposure barely reduced the viability of neither planktonic cultures nor biofilms of any strains. On the contrary, for all selected strains, addition of vancomycin at T0=6h followed by a 24 hours exposure significantly and more efficiently reduced the viability of planktonic cultures than that of biofilms, which was not decreased at all in the case of strains of ribotype WRT 005, 009 and 035. The biofilms of these selected strains of eight ribotypes, compared to their planktonic cultures, therefore showed a higher tolerance to vancomycin, suggesting that they could play a role in persistence.

## 1. Introduction

*Clostridioides difficile* is a Gram-positive, spore-forming obligate anaerobe and an opportunistic entero-pathogen of humans and animals, whose spores are ubiquitous in the environment (Lim *et al*., 2020). It is widely distributed among various animals, including wild animals, farm animals like livestock and pigs, pets and horses (Weese, 2020). As a human pathogen, *C. difficile* is considered as an “urgent threat” by the Centers for Disease Control and Prevention in USA since 2013 and this status has been confirmed in 2019. *C. difficile* is the leading cause of healthcare-associated infections in the United States and of post-antibiotic gastrointestinal illnesses. Indeed, previous antibiotic treatments are among the main risk factors for developing a *C. difficile* infection (CDI) (Taggart *et al*., 2021). Moreover, CDI are often recurring, by approximately 20% of cases the first time and by a more and more increasing rate after each episode (Tresman and Goldenberg, 2018). Re-infection by the same strain after the end of antibiotic treatment (fidaxomicin, vancomycin) has been described (Figueroa *et al.,* 2012), suggesting both poor efficiency of treatment and strain persistence. Finally, CDI is also becoming a common community-acquired infection, affecting younger and otherwise healthy people (Guery *et al*., 2019; De Roo and Regenbogen., 2020).

The *C. difficile* infection cycle in humans and animals begins by oral contamination by hospital or environmental spores (Smits *et al*., 2016). The use of broad-spectrum antibiotics alters the gut microbiota composition and metabolism, thus promoting *C. difficile* spore germination and emergence of vegetative cells, which can grow and express their virulence, finally leading to disease (Smits *et al*., 2016). Virulence is primarily mediated by toxins. The two major ones are toxin A (TcdA) and toxin B (TcdB), even though TcdB alone is sufficient to cause disease, while the binary toxin (CDT) is considered to have an accessory role. Finally, *C. difficile* cycle ends by sporulation and dissemination of spores in the environment through the faecal route. Spores are highly resistant to many stresses, including oxygen, heat stress and disinfectants: they are able to persist in the environment and to be transmitted to another host (Dawson *et al*., 2011).

Biofilms are sessile communities living within self-produced extracellular matrices (Hall-Stoodley and Stoodley, 2009). In the case of pathogen species, biofilms can be responsible for chronic infections characterised by the persistence and increased tolerance to antibiotics of the pathogen (Bjarnsholt., 2013; Lebeaux *et al*., 2014).

*C. difficile* has been shown to form biofilms *in vitro* on different abiotic surfaces and in different culture systems using different media (Dapa *et al*. 2013; Dapa and Unnikrishnan., 2013; Maldarelli et *al*., 2016; Pantaléon *et al*., 2014; Poquet *et al*., 2018).

The structures of *in vitro* biofilms are diverse and can include multicellular aggregates as well as mats, which have also been observed *in vivo* (Maldarelli et *al*., 2016; Poquet *et al*., 2018; Soavelomandroso *et al*., 2017, Dawson *et al*., 2021). Their matrix is made of extracellular DNA, proteins and polysaccharides (Kavanaugh *et al*., 2019). Some biofilms have been shown to express toxin genes (Dapa *et al*., 2013; Dubois *et al*., 2019; Semenyuk et *al*., 2014), and to display low susceptibility to antibiotic treatment (Dapa et *al*., 2013; Semenyuk *et al*., 2014). In addition, late biofilms are able to produce spores, suggesting that they could contribute to spread and persistence (Dapa *et al*., 2013; Dawson *et al*., 2012; Dubois *et al*., 2019; Semenyuk *et al*., 2014; Tijerina-Rodríguez *et al*., 2019). Finally, on the basis of these *in vitro* data, *C. difficile* biofilms could play different roles *in vivo*: virulence, protection against treatments, tolerance, persistence, dissemination, and possibly re-infections.

Early studies on *C. difficile* biofilms focussed on model strains, like 630 or R20291, which were initially isolated as clinical isolates, or like 630Δ*erm*, which had been obtained as a erythromycin-sensitive mutant of strain 630 (Hussain *et al.,* 2005). During their history of laboratory growth, model strains acn accumulate spontaneous mutations changing their phenotypes (Collery *et al.,* 2016). In *Bacillus subtilis,* a Gram-positive spore-forming species, in the 2000-2010 period, non-domesticated strains compared to laboratory strains were shown to display unique multicellular phenotypes, including biofilms of specific structures (Aguilar et *al*., 2008; Bridier et *al*., 2011). Consequently, in *C. difficile*, natural isolates and isolate collections are raising more and more attention (Ais et *al*., 2023; Morais *et al*., 2022; Pantaléon et *al*., 2018; Rahmoun *et al*., 2021; Tijerina-Rodriguez et *al*., 2019; Wultanska et *al*., 2023).

Here, we took advantage of our CloDifEqui collection of *C. difficile* undomesticated strains isolated from twenty-six equids necropsied in Normandy, France, between 2018 and 2021 (Petry *et al*., 2024). We chose one strain per animal, the one recovered from animal caecum when available, but included the two strains of different ribotypes isolated from the same co-colonized animal, finally leading to twenty-seven strains representative of *C. difficile* molecular diversity. Twenty strains belonging to eight ribotypes were toxigenic and able to produce from one to three toxins: they were asymptomatically carried by sixteen equids and caused CDI in the four remaining ones. Seven non-toxigenic strains of three ribotypes were carried, as expected (Petry *et al*., 2024). In this study, we aimed to evaluate the ability of undomesticated, diverse strains of *C. difficile* to form biofilms *in vitro* and to characterise their morphology and properties.

## 2. Materials and Methods

### 2.1. Bacterial strains and general growth conditions

27 representative members of CloDifEqui, the collection of *C. difficile* strains previously isolated from equids (Petry *et al*., 2024) were chosen (Table 1). The laboratory model strains 630Δ*erm* and VIP 10463 were used as controls when indicated.

**Table 1.**
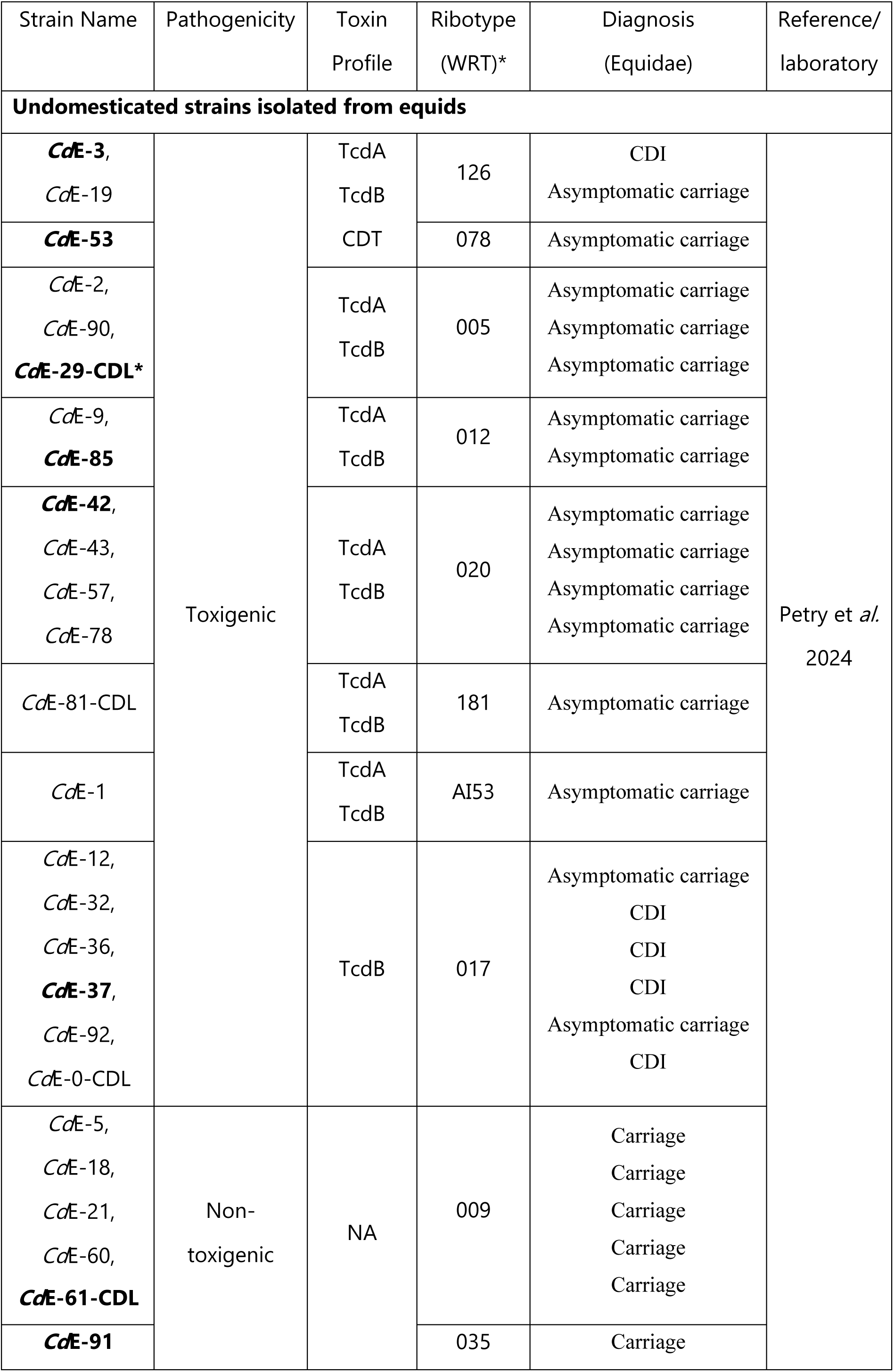
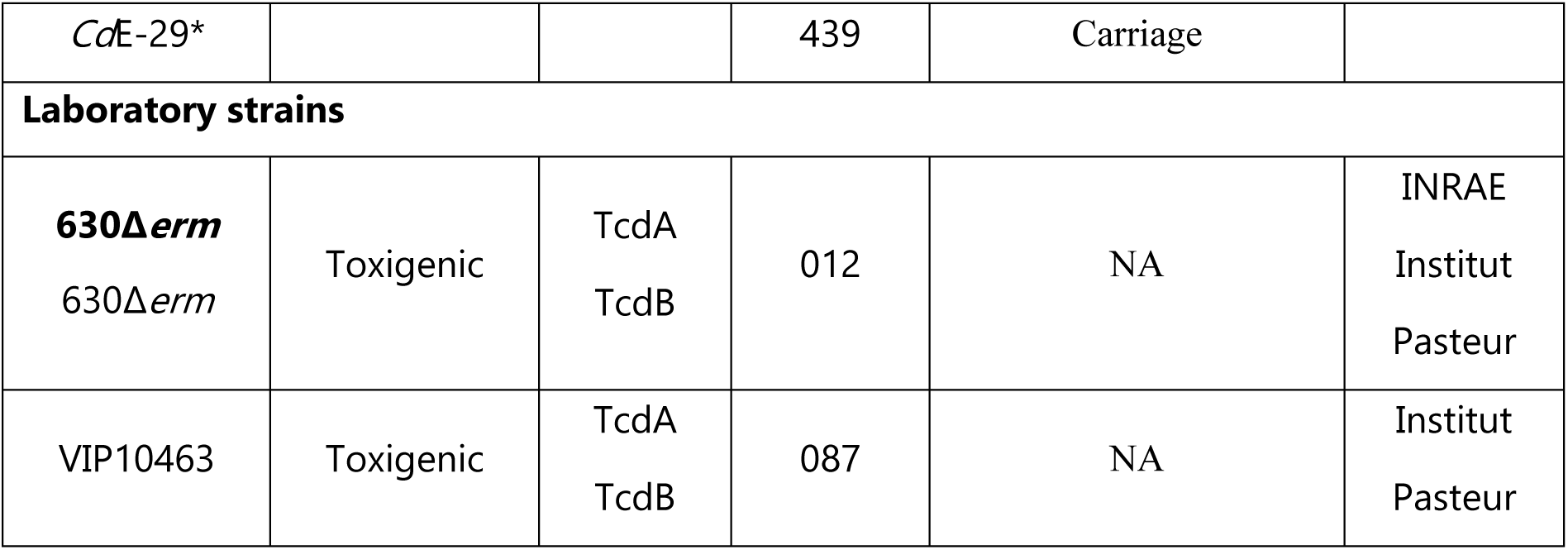
*C. difficile* strains used in this study. Undomesticated strains (CloDifEqui collection) had been isolated from Equids necropsied in Normandie (Petry *et al*. 2024). Laboratory strains were from laboratory collections. Strain ribotype (PCR-ribotype assigned according to the WEBRIBO database), toxin profile and pathogenicity are indicated, together with diagnosis of CDI, asymptomatic carriage (toxigenic strain) or carriage (non- toxigenic strain) in the case of equine strains (Petry *et al*., 2024). Representative strains selected for a more detailed study (see below) are indicated in bold. NA not applicable. * is indicated for the two strains that had been isolated from the same animal (#29), either from its caecum or from another watery intestinal segment (CDL) (Petry *et al*., 2024).

For routine experiments, strains were isolated from a frozen stock at -80°C on ChromID agar plates (BioMérieux, France), and a single colony was streaked on Brain Heart Infusion (BHI) agar plates. A single colony was inoculated in 10 mL BHIS glucose medium (brain-heart infusion supplemented with yeast extract, 0.1% cysteine, 0.1 M glucose) (Becton Dickinson, France; Merck, Germany) in Falcon tubes for overnight growth. All incubations were performed under anaerobic conditions (90% N2, 5% CO2 and 5% H2) at 37°C in a Freter cabinet.

### 2.2. Biofilms and planktonic cultures conditions

Biofilms were grown in 96-well microtiter plates as previously described, except in BHIS glucose (Poquet et *al*., 2018) and planktonic cultures were grown under the closest possible conditions. Briefly, overnight cultures of all strains were first diluted in BHIS glucose to an OD600nm of 0.3. 250 µL of these bacterial suspensions at OD600nm 0.3 were inoculated either into a 96-well polystyrene plate (Greiner, France), for biofilms, or into Eppendorf tubes, for planktonic cultures. For biofilms, the plate outermost wells were filled-in with PBS to prevent evaporation during incubation at 37°C. A primary incubation of 2 hours was performed to allow cell adhesion, while in parallel, planktonic cultures were vigorously vortexed every 30 minutes to prevent adhesion. After the 2h- incubation, most of the liquid was carefully removed by pipetting and 250 µL of fresh BHIS glucose medium were added. Biofilms and planktonic cultures were then let to grow from initially adhesive cells and non-adhesive cells respectively, by incubation for 48 hours. During this secondary incubation, planktonic cultures were again vigorously vortexed every 30 minutes except overnight and finally once again at the end of incubation, before recovery or treatment. After 48h-growth, biofilm pH was measured using pH paper (Merck, Germany).

### 2.3. Crystal violet (CV) staining of biofilms Biomass of either adhesive biofilm or both adhesive and wash-resistant biofilm

In the literature (Dawson *et al*., 2012; Dawson *et al*., 2021; Dapa *et al*., 2013; Pantaléon et *al*., 2015; Pantaléon *et al*., 2018; Purcell at *al.*, 2017), the biomass of *C. difficile* biofilms formed at the bottom of wells in a microtiter plate are usually quantified by CV staining after two treatments: i) removing the upper liquid phase (assumed to contain planktonic cells) by turning the plate upside down, followed by ii) biofilm washing. We observed that biofilms were fragile and partially removed by washing and decided to evaluate the effect of biofilm washing or not before staining. In the following, the biomass values were measured for the biofilm just after turning the microplate upside down (adhesive biofilm) or after both turning the microplate upside down and washing of the adhesive biofilm (adhesive and wash-resistant biofilm). Biofilm washing was performed twice with 100 µL of sterile 1x PBS. Biofilms were then fixed with 100 µL of a mixture of 75% ethanol and 25% acetic acid (v/v) for 20 minutes, dried at 37°C for 20 minutes, stained with 250 µL of 0.2% crystal violet (Merck-Sigma, Germany) and incubated at 37°C for 30 minutes. After two steps of gentle washing with 300 µL of sterile 1x PBS, crystal violet from stained biofilms was extracted with 250 µL of ethanol and its absorbance at 570 nm was measured. Background was quantified in two wells with BHIS glucose medium and subtracted from any biofilm measure. The biomass of each undomesticated strain biofilm was normalised using that of 630Δ*erm* biofilm present in the same plate (assigned to 100%). The experiment was performed in six independent replicates for each equine strain, leading to a total of forty-eight replicates for the laboratory strain used as a reference, 630Δ*erm*.

### 2.4. Confocal Laser Scanning Microscopy (CLSM): intact biofilm observation

Biofilms were observed by CLSM, essentially as described by Poquet et *al* (2018). Live-dead staining (Filmtracer™ LIVE/DEAD^TM^ Biofilm Viability Kit, Thermo Fisher Scientific, United States) was performed by carefully adding a mix of Syto9 (targeting DNA of live cells) and propidium iodide to reach final concentrations of respectively 9.5 µM and 57.1 µM onto the biofilm (without perturbing it) and by incubating for 2 hours at 37°C under anaerobiosis (using AnaeroGen, Oxoid, Biomérieux, France).

The 3D structure of intact biofilms was studied *in situ* by CLSM using a Leica SP8 AOBS inverted laser scanning confocal microscope (Leica Microsystems, Germany) and a ×63 water immersion objective (N.A. = 1.2) on INRAE MIMA2 core facility (https://mima2.jouy.hub.inrae.fr/). For each strain, the biofilms of three independent clones were analysed and at least two series of stacks were randomly acquired in each well. Stacks of horizontal images (z stacks of xy images) were sequentially acquired in 1 µm increments with a scanning speed of 600 Hz. Fluorescence excitation was achieved using a 488 nm argon laser line set at 30% intensity *via* the argon potentiometer and 10% via the AOTF (Acousto-Optic Tunable Filter). The emitted fluorescence was recorded by the AOBS system using two simultaneous photomultiplier tubes (PMTs): between 497 and 570 nm for the green fluorescence of SYTO^9^ and between 637 and 715 nm for the red fluorescence emitted by propidium iodide.

Observations and recordings were made on three independent biofilms for each equine strain (all grown from independent clones during two to three different experiments) and twelve independent biofilms for the reference strain. Each biofilm was observed in duplicate.

### 2.5. Analysis of CLSM image

#### Three-dimensional images

z-stacks were acquired during CLSM experiments and analysed using BiofilmQ software (version 1.0.1) to quantify geometric parameters of biofilm structure, including total biovolume, mean thickness and surface roughness (Hartmann *et al*., 2021). Image processing was standardised across all fluorescence channels. First, images were denoised using convolution (dxy = 5 and dz = 3) and then segmented into two classes using the OTSU thresholding method with a sensitivity of 2. The detected signal was downscaled to 3.68 µm^3^ cubes and small objects were removed using a threshold of 0.5 µm³ to eliminate residual noise. In each experiment, the data from each strain were normalised using those of the reference strain 630Δ*erm* present on the same plate. A PHP script (version 7.4) was used to aggregate the values obtained from each experiment in a spreadsheet.

#### Two-dimensional images

he z-stacks were imported into ImageJ software (version 1.54j). The appropriate colour was assigned to each fluorescence channel: green (Syto^9^), and red (propidium iodide). Merging could lead to an orange colour for objects stained by both Syto^9^ and propidium iodide fluorescences. xy images were extracted close to the polystyrene surface at 16<z<29.

### 2.6. Sporulation assay

After 48 hours of growth, the biofilms and planktonic cultures of selected strains were harvested and divided into two aliquots, one of which was exposed to 65°C during 25 min (Dawson et *al*., 2012) to kill vegetative cells and specifically quantify spores. Each aliquot was then serially diluted (by a 10-fold factor) into sterile 1x PBS. All dilutions were plated on BHIS agar plates supplemented with 0.1% taurocholate to promote germination. For each dilution, colony forming units (CFU) were counted after 24 hours of growth at 37°C under anaerobic conditions and CFU/mL calculated. Sporulation rate was the percentage of spores after heat treatment compared to the total cell population, as determined in the absence of treatment. Four independent biological replicates were performed.

### 2.7. Vancomycin tolerance assay

We compared the tolerance of selected strains to vancomycin after growth in biofilm or planktonic cultures under the same conditions, in BHIS glucose medium for 48h. The minimum inhibitory concentration (MIC) of each CloDifEqui strain had previously been determined (Petry *et al*., 2024), and notably that of each selected strain (CdE-91: 0.125 mg/L, CdE-53 and CdE-85: 0.5 mg/L, CdE-37 and CdE-42: 0.75 mg/L and CdE-3: 1 mg/L). As a preliminary experiment, the MIC value of reference strain 630Δ*erm* was established by the same method, using Etest (bioMérieux, France) on blood agar plates (BBA, bioMérieux, France), and found to be of 0.75 mg/L. Based on all these data, a concentration of 8 MIC, ranging from 1 mg/L to 8 mg/L depending on the strain, was used.

After an initial growth of 6 or 24 hours (T0), the biofilms and planktonic cultures of each strain were enumerated and treated or not with vancomycin at 8 MIC. They were then further incubated for 24 hours. After vancomycin exposure or not for 24 hours (Tf), planktonic cultures and biofilms were recovered by pipetting. After addition of 100 µL of sterile 1x PBS and vigorous vortexing, serial dilutions (by a 10-fold factor) in 1x PBS were plated on BHIS agar with 0.1% taurocholate. Finally, CFU were counted as previously described for the sporulation assay. Experiments were performed in 4 independent replicates.

### 2.8. Toxins detection (TcdA and TcdB)

The biofilms were harvested by gently dislodging them with a micropipette and transferring them to an Eppendorf tube. A volume of 100 µL of PBS was added to both the biofilms and the planktonic cultures, which were then frozen at -20°C. Microtiter plates (Nunc Maxisorp, BioLegend, United States) were coated with anti-TcdB or anti-TcdA capture antibodies (LsBio, United States) at a concentration of 1 µg/mL and incubated overnight at 4°C. The plates were washed three times with 200 µL of 1x PBS containing 0.05% Tween 20. Protein binding sites were blocked with 200 µL of blocking buffer (SuperBlock Blocking Buffer in PBS, Thermo Fisher Scientific, United States) for 1 hour at room temperature, followed by three additional washes with 200 µL of 1x PBS and 0.05% Tween 20. Biofilms and planktonic cultures were added and incubated for 1 hour at room temperature, followed by three washes. Purified toxins A at 0.1 µg/mL and B at 0.2 µg/mL were used as standards, with 100 µL of each toxin added to each well and incubated for 1 hour at 37°C. After washing, 100 µL of biotinylated anti-toxin B antibody (LsBio, United States) was added, followed by high sensitivity streptavidin-HRP conjugate (Thermo Fisher Scientific, United States) or HRP-conjugated anti-toxin A antibody (LsBio, United States). The antibody signal was detected using TMB substrate (Thermo Fisher Scientific, United States) at 450 nm on an ELISA plate reader. Two laboratory strains were included as positive controls: 630Δ*erm* (for TcdA and TcdB) and VIP10453 (for TcdB) [Table 1]. The experiment was performed in 2 independent replicates

### 2.9. Graphical Representation of Data and Statistical Analysis

Statistical analyses were performed using the non-parametric Wilcoxon-Mann- Withney test (for sample sizes of less than 30) with GraphPad Prism software (version 8.0.2). Significance threshold was p < 0.05.

## 3. Results

### 3.1. Evaluation of Biomass and Adherence of *C. difficile* Biofilms

We first examined twenty-seven non-domesticated strains together with the reference strain 630Δ*erm* for their ability to form biofilms, using a standard method allowing quantification but destructing biofilm structure. Biofilms were obtained after removing the liquid upper phase (adhesive biofilm) or after both removing the liquid upper phase and washing biofilm cells (adhesive and wash-resistant biofilm), and finally stained by crystal violet (CV) to quantify their biomass. The results for each strain were normalised by comparison to those of the reference strain 630Δ*erm*.

We first evaluated the biomass of adhesive biofilms. Compared to 630Δ*erm*, eleven and eight strains formed adhesive biofilms of a significantly higher or lower biomass, respectively, while the adhesive biofilms of the remaining eight strains showed a similar biomass [Figure 1A]. When strains of the same ribotype were considered as a whole, biomass was significantly higher for the mean adhesive biofilm of ribotype WRT009 strains than for the mean adhesive biofilm of ribotypes WRT012 (including the reference 630Δ*erm*) or WRT017 [Figure 1B]. As a whole, non-toxigenic strains, compared to toxigenic ones, appeared to form a mean adhesive biofilm of higher biomass [Figure 1C]. In the case of toxigenic strains, we also analysed the results with respect to strain phenotypes in equids, i.e either *in vivo* virulence/CDI or asymptomatic carriage [Table 1, Petry et *al*., 2024] and found no statistically significant difference (not shown).

**Figure 1.**
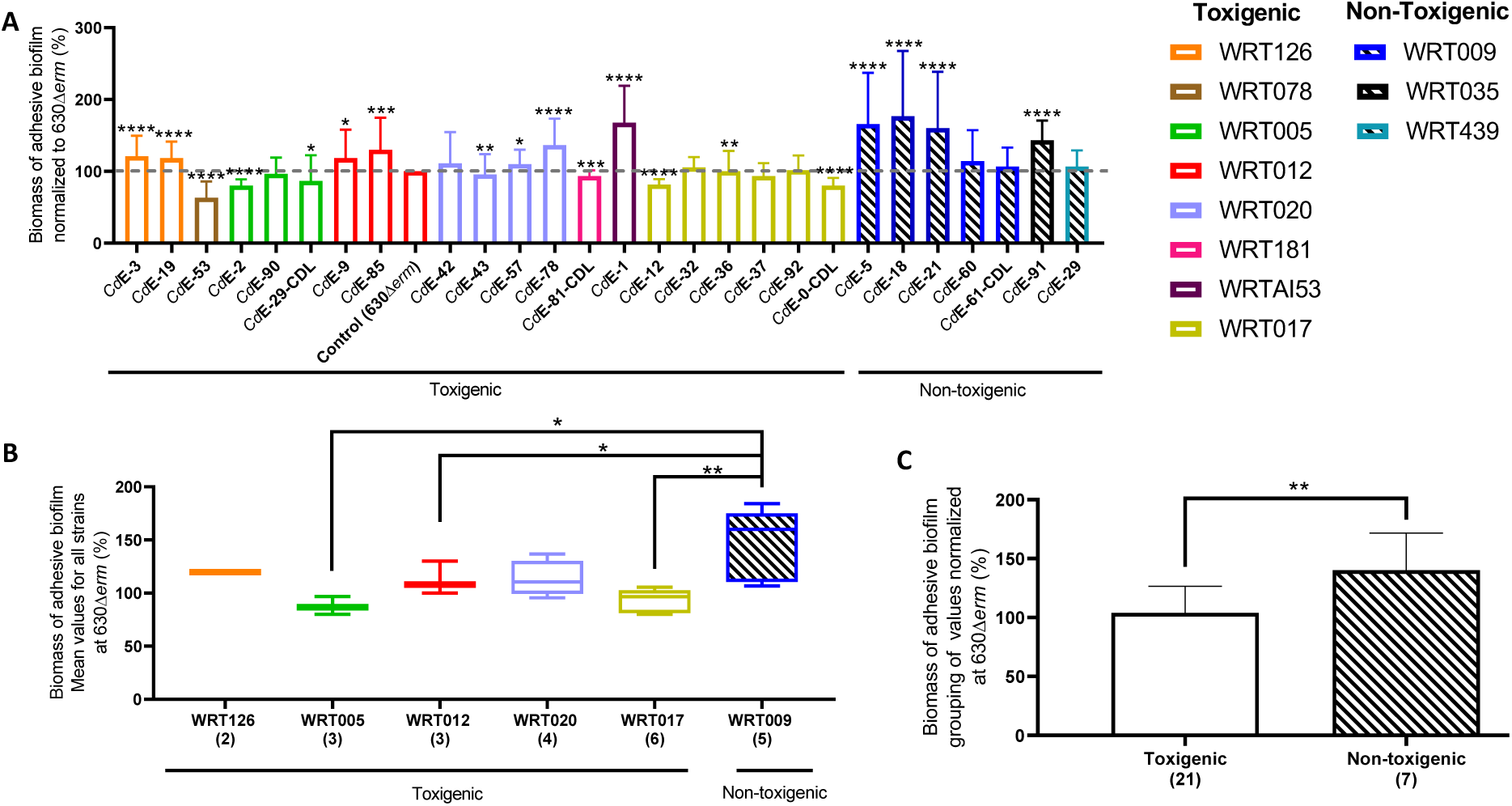
: Biomass of adhesive biofilm, as revealed by CV staining. Twenty-seven undomesticated strains of *C. difficile* (Table 1) were studied together with the laboratory strain 630Δ*erm* used as a reference. After having removed the upper liquid phase, the biomass of adhesive biofilm was quantified by CV staining. Values were the mean of six measures for any undomesticated strain and forty-five measures for the reference strain 630Δ*erm*. The result for each equine strain was normalised with respect to the value obtained for the reference strain 630Δ*erm* present on the same plate and assigned to 100%. (A) The biomass of each strain adhesive biofilm, expressed as a percentage of the biomass of 630Δ*erm* adhesive biofilm (a red dotted baseline is drawn to indicate 100%). (B) The biomass of the mean adhesive biofilm of any ribotype containing at least two strains. (C) The biomass of the mean adhesive biofilm of all toxigenic strains and of all non-toxigenic strains. A colour was assigned to each ribotype (see inset at the top in the right). Wilcoxon-Mann-Whitney test * p<0.05, ** p<0.01, *** p<0.001, **** p<0.0001.

We then also measured the biomass of the adhesive and wash-resistant biofilm [Figure 2], as performed in most studies (Dawson *et al*., 2012; Dawson *et al*., 2021; Dapa *et al*., 2013; Pantaléon *et al*., 2015; Pantaléon *et al*., 2018; Purcell at *al.*, 2017). Compared to 630Δ*erm*, seventeen strains, including all five strains of WRT 009 and all four strains of WRT 020, displayed biofilms of significantly higher biomass, indicating their higher resistance to wash [Figure 2A]. When strains of a given ribotype were considered as a whole, the mean biofilm of ribotype WRT009 was significantly more resistant to wash than that of WRT012, WRT005 and WRT017, and the same held true for WRT020 compared to WRT017[Figure 2B]. Similarly, all non-toxigenic strains displayed a mean biofilm more resistant to wash than that of all toxigenic strains [Figure 2C]. There was no difference between toxigenic strains according to their phenotypes in equids (Table 1, Petry et *al*, 2024) (not shown).

**Figure 2.**
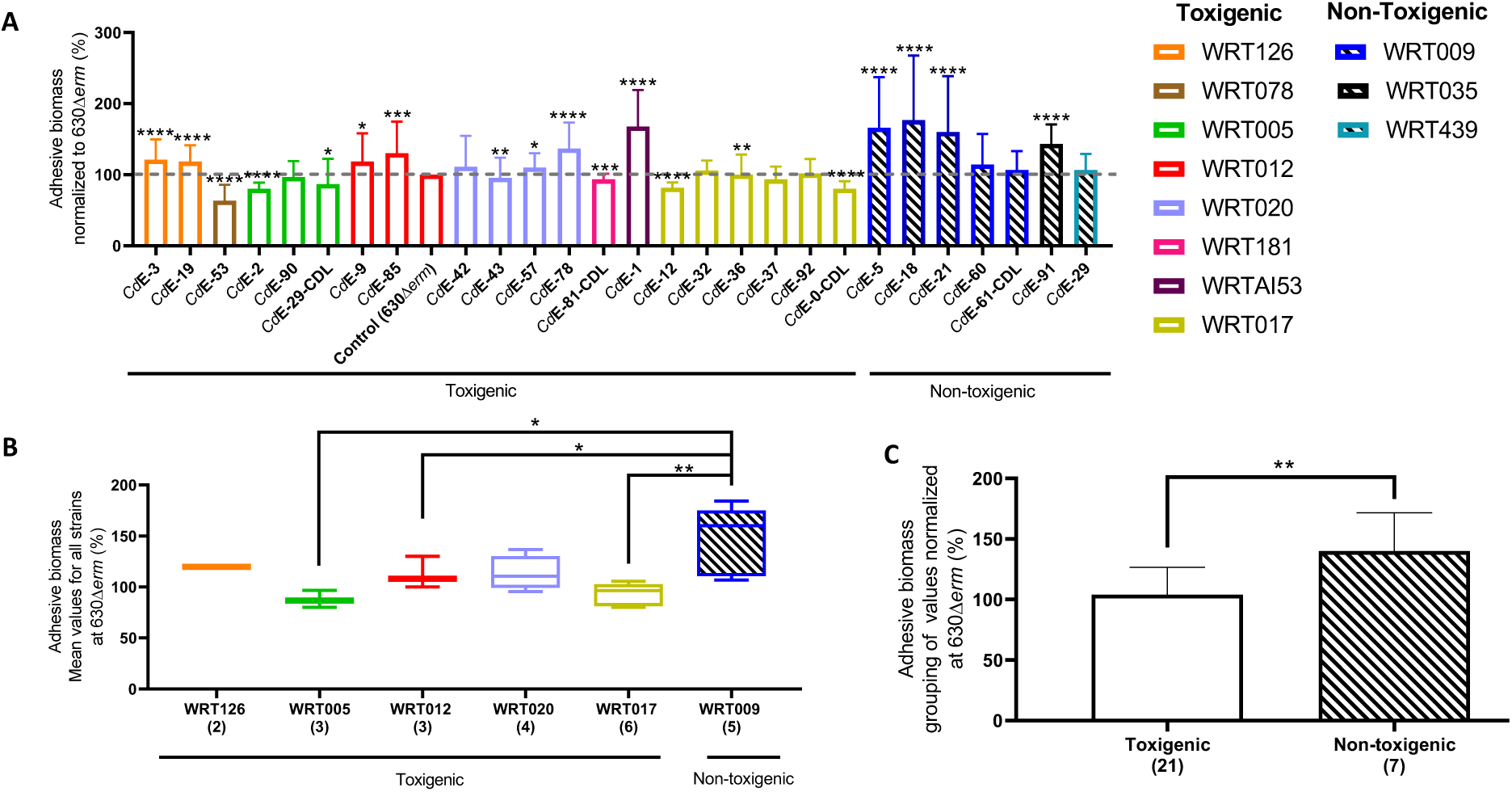
: Biomass of both adhesive and wash-resistant biofilms. Here, CV staining was performed after both removing the upper liquid phase and washing. Otherwise, all is like in Figure 1 (see Figure 1 legend).

### 3.2. The 3D structure of intact biofilms observed *in situ*

We also studied intact biofilms, without any perturbation, by CLSM after live/dead staining, to get access to their 3D architecture *in situ* (Poquet et *al*., 2018). The biofilms of all strains [Supplementary data 1] and in particular those of eight selected strains belonging to the most common ribotypes [Figure 3] were observed. Three parameters describing biofilm structure (volume, thickness, roughness) were measured [Figure 4, Supplementary data 3].

**Figure 3.**
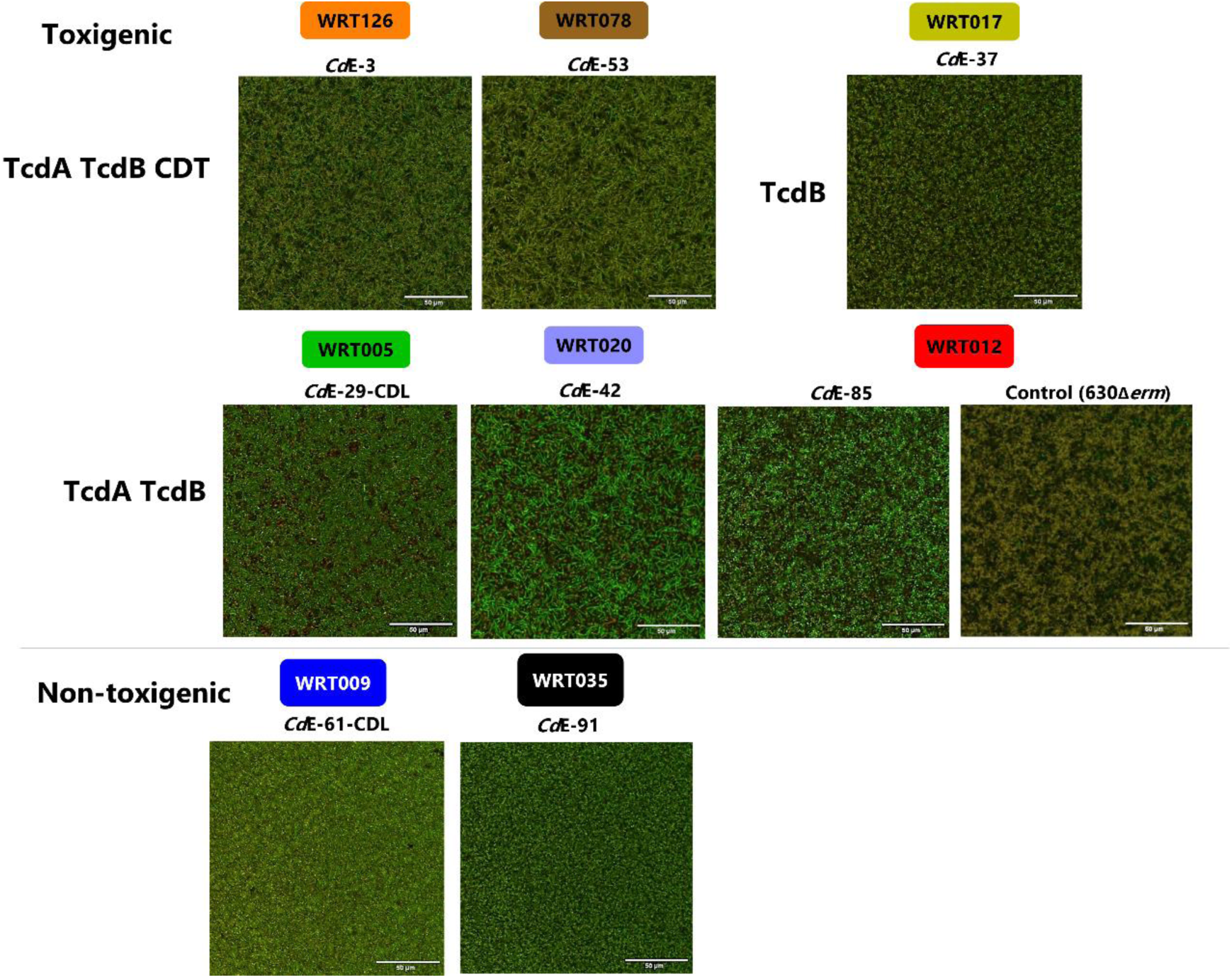
: The structure of intact biofilms observed *in situ* by CLMS after live-dead staining. For one representative isolate of the eight most common ribotypes found in equids (strain name and ribotypes (WRT) are indicated) together with the reference strain 630Δ*erm*, a representative xy image was chosen at the biofilm basement (at a z position (height) comprised between z=17nm and z=28nm corresponding to the bottom of biofilms).

**Figure 4.**
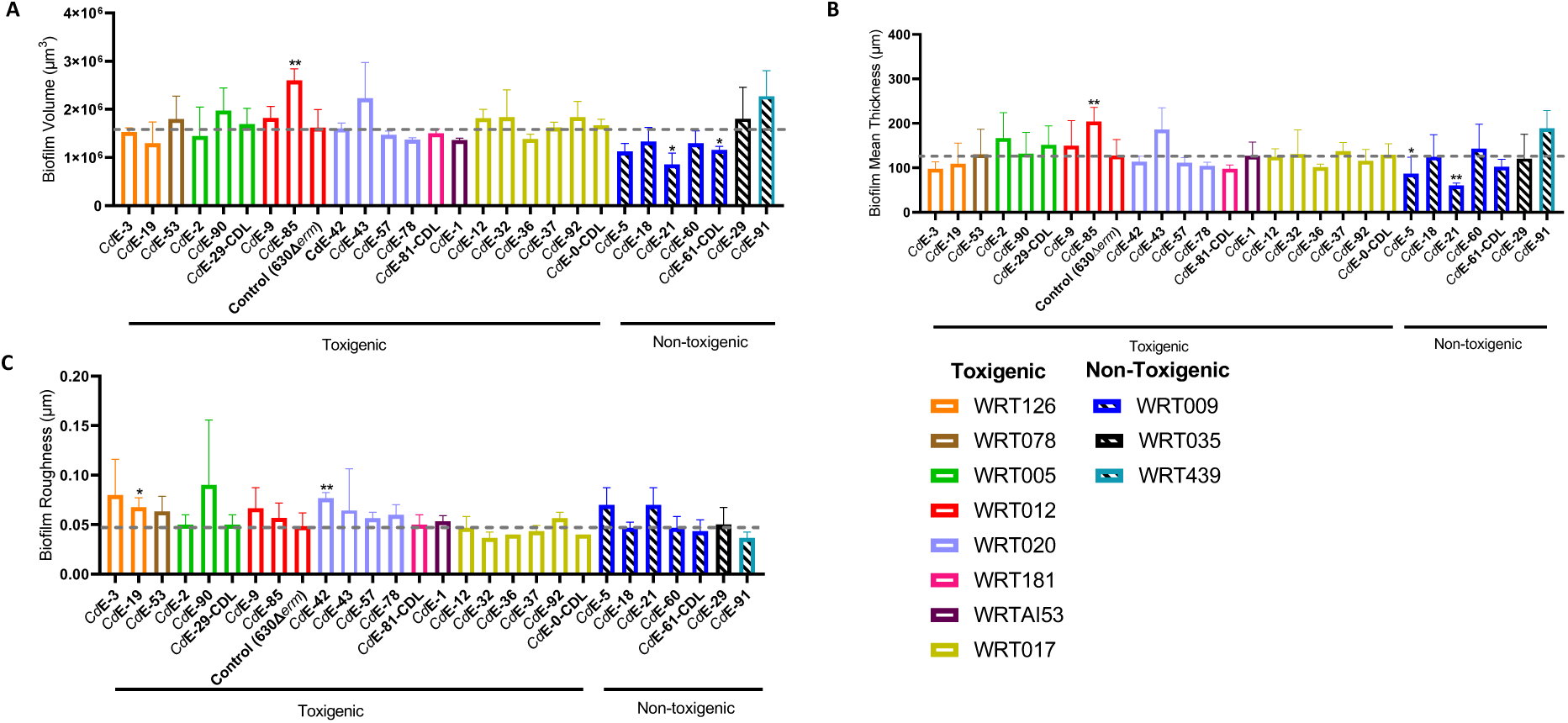
: The 3D structure of intact biofilms analysed according to strain ribotype. After live dead staining and observation of intact biofilms by CSLM, their microscopic 3D structure was evaluated using BiofilmQ software. The experiment was repeated 3 times for each equine strain. (A) Biofilm biovolume. (B) Biofilm mean thickness. (C) Biofilm roughness, which is defined as the standard deviation of thickness measured for vertical sections along each biofilm. Wilcoxon-Mann-Whitney test * p<0.05, ** p<0.01, *** p<0.001.

Compared to the control strain 630Δ*erm*, all strains formed biofilms with similar structures. For each strain, xy sections representative of the biofilm structure close to the polystyrene surface of the plate well were observed [Figure 3, Supplementary data 1]. No major difference could be evidenced between strains, even though the biofilm cells of WRT009 strains [Supplementary data 3] and in particular of *Cd*E-61-CDL [Figure 3], seemed to be slightly more tightly packed than those of other strains.

Compared to that of the control strain, the biofilm of strain CdE-85 was characterised by a significantly higher volume and thickness, while the opposite hold true for the biofilm of strain CdE-21 [Figure 4A and 4B]. The biofilm of strain CdE-61-CDL had a smaller volume [Figure 4A]. The biofilms of strains CdE-19 and CdE-42 were characterised by a higher roughness [Figure 4C].

We then analysed the strains of the same ribotype as a whole [Supplementary Data 2]. The mean biofilm of WRT009 strains displayed a significantly smaller volume than the mean biofilm of any of the following ribotypes: WRT005, WRT012, WRT020 or WRT017 [Supplementary Data 2A].

### 3.3. Comparison of sporulation and toxin production in biofilms and planktonic cultures

We compared the ability of biofilm and planktonic cells to produce spores after 48 hours of growth in BHIS glucose medium for one selected strain of each common ribotype [Figure 6]. The percentage of sporulation was similar between planktonic cultures and biofilms for all strains (between 5 % and 10% in average) except three: in strains CdE-37, CdE-29-CDL and CdE-91, spore production was significantly lower in biofilms (0.3% to 2%) compared than in planktonic cultures [Figure 5A]. We observed no difference between toxigenic and non-toxigenic strains [Supplementary Data 4].

**Figure 5:**
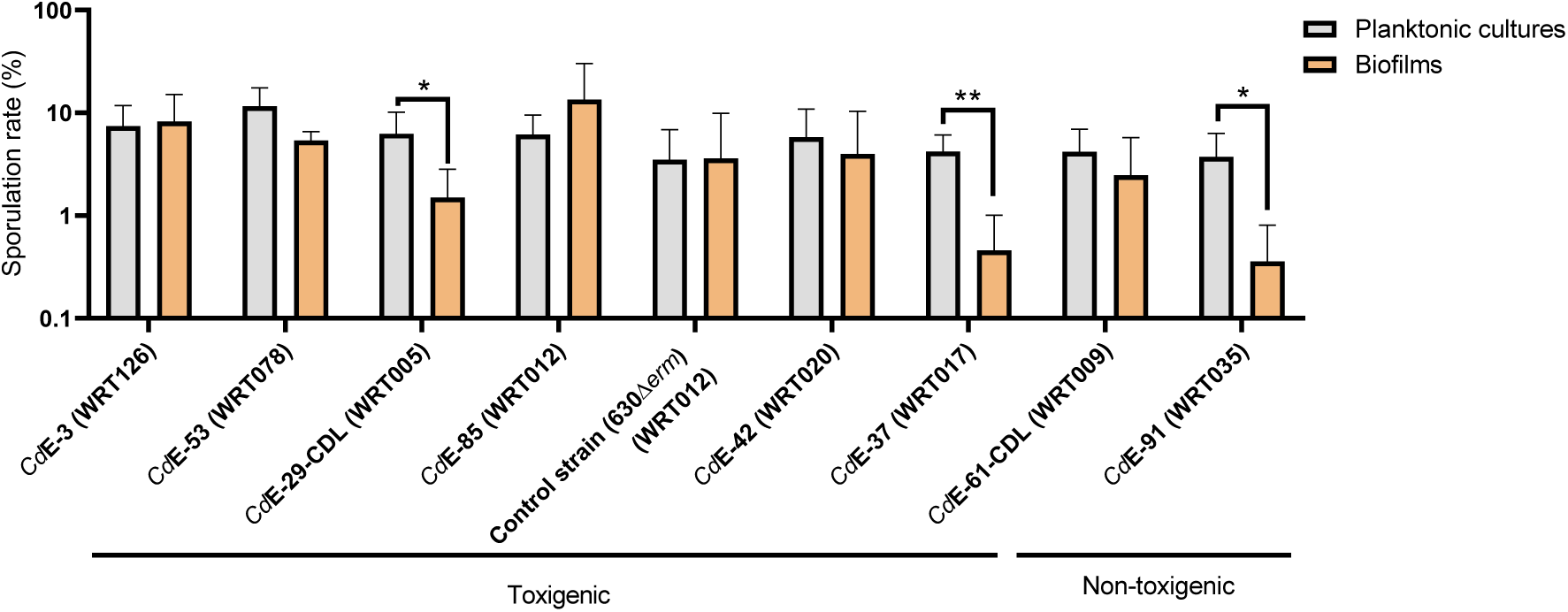
Sporulation rate (percentage of heat-resistant spores compared to the total cell population) of planktonic cultures and biofilms after growth for 48h in BHIS glucose medium. Eight strains representative of the most common ribotypes in equids (the same as in Figure 3) were studied. Results of cfu counting are shown using a logarithmic scale. The experiment was repeated four times for each strain. The pH of the BHIS glucose starting medium was basic (∼8.0) and that of planktonic cultures and biofilms after 48h in this medium was acidic (between 5.0 and 6.5), indicating growth and fermentative activity of *C. difficile* cells. Wilcoxon-Mann-Whitney test: * p < 0.05, ** p < 0.01.

**Figure 6:**
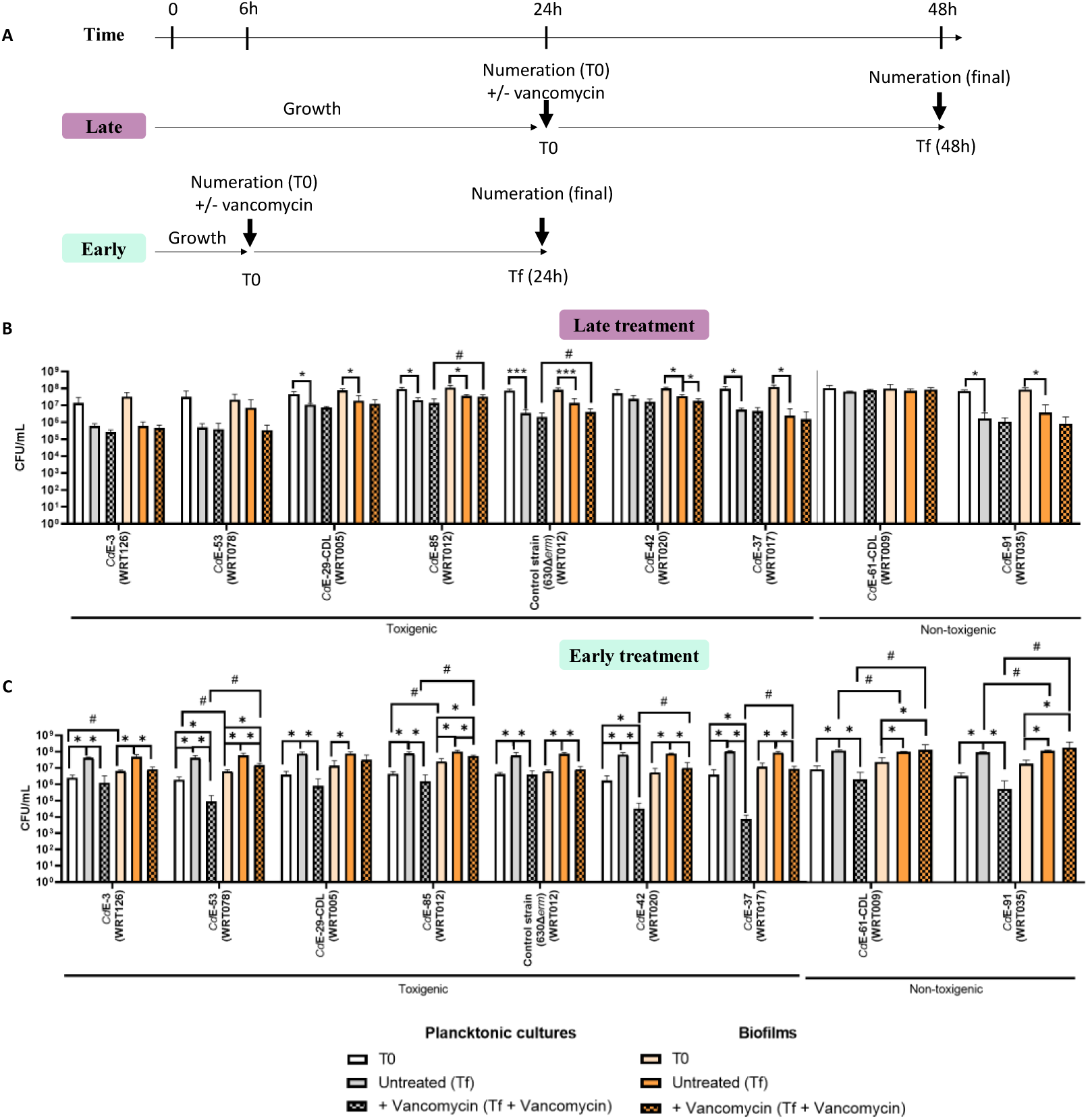
Tolerance of biofilms and planktonic cultures to vancomycin. The same strains as in Figure 3 and 5 were studied. For each of them, vancomycin was added or not at a concentration of 8 MIC (Petry et *al*. 2024). (A) Summary of experimental conditions. Vancomycin was added at T0, after either 24 hours (late treatment) or 6 hours of growth (early treatment). After vancomycin addition or not, cell incubation went on for further 24hours, until Tf. Cfu counting was performed at T0 (before addition) and at Tf (in the presence of vancomycin or not). The experiment was repeated four times for each strain, and mean cfu/mL calculated (B) Late addition of vancomycin (T0 = 24h). (C) Early addition of vancomycin (T0 = 6h). In both (B) and (C), planktonic cultures are shown as white bars at T0 and as grey bars at Tf, and biofilms as pale orange bars at T0 and dark orange bars at Tf. Vancomycin exposure is indicated by hatched bars. Cfu/mL are shown using a logarithmic scale. Wilcoxon-Mann-Whitney test: * p < 0.05, ** p < 0.01, *** p < 0.001, for the comparison between vancomycin exposure or not for each strain and each growth mode (either planktonic culture or biofilm) and strain. # p < 0.05 for the comparison for each strain between planktonic culture and biofilm, both under the same condition of vancomycin exposure or not.

We also tested, for each selected strain, the ability of biofilm and planktonic cells to produce TcdA and TcdB toxins by Elisa. As expected, we were able to detect toxins in two positive controls: purified TcdA and TcdB proteins and toxins produced by the reference strain 630Δ*erm* under known permissive conditions, after overnight growth in TY medium (Antunes et *al*., 2011). On the contrary, neither TcdA (data not shown) nor TcdB [Supplementary Data 5] could be detected for any equine strain, after 48 hours of either biofilm or planktonic growth in BHIS glucose medium.

### 3.4. Tolerance to vancomycin of biofilms and planktonic cultures

We finally studied the biofilms and planktonic cultures of each selected strain for their tolerance to vancomycin, an antibiotic widely used to treat CDI [Figure 7]. Vancomycin was added or not at a concentration of 8 MIC for each strain (Petry et *al*., 2024).

We studied the effect of vancomycin added after 24h of growth and for a 24 hours period (‘late treatment’, Figure 7A). We compared the final vegetative cell counts after vancomycin exposure or not. Despite the use of vancomycin at a highly inhibitory concentration, there was no significant difference for any strain, neither in planktonic culture nor in biofilm [Figure 7B]. We noticed that in the absence of treatment, cells at the end of the experiment (at Tf) compared to the beginning (at T0) had not multiplied for any strain and any growth mode (planktonic culture or biofilm). The absence of cell growth between Tf and T0, as observed for untreated cells, might hide any vancomycin effect and contribute to its lack of efficiency on treated cells.

We therefore decided to add vancomycin much earlier, after 6 hours growth, without modifying exposure time for 24 hours (‘early treatment’, Figure 7A). Under these conditions of early addition, as expected, cell counts of untreated planktonic cultures and untreated biofilms at Tf compared to T0 significantly increased for all strains [Figure 7C]. Moreover, in vancomycin-treated compared to untreated planktonic cultures of all studied strains, the final cell counts significantly decreased, by a 100- to 1000-fold factor depending on the strain [Figure 7C]. Vancomycin addition in early biofilms resulted in a significant but weaker decrease of the final cell counts, by only a 10-fold factor [Figure 7C]. These results showed that, in the case of strains of eight different ribotypes, vancomycin at a highly inhibitory concentration was barely or not efficient when added lately, and more efficient when added early. Moreover, for these eight strains, even after early addition, vancomycin was much more efficient against planktonic cells than against biofilm cells grown under the same conditions, demonstrating the higher vancomycin tolerance of the latter compared to the former.

## 4. Discussion

Here, we addressed the question of whether different *C. difficile* strains could form biofilm of different morphologies or properties, using a collection of non-domesticated *C. difficile* strains, which had been isolated from equids and belonged to eleven ribotypes. Our twenty-seven non-domesticated strains were all able to form biofilms after 48 hours of growth in rich BHIS glucose medium. Several studies previously showed the ability of several *C. difficile* laboratory strains to form biofilms *in vitro* (Dapa *et al*., 2013; Dawson *et al*., 2012, Maldarelli et *al*., 2016, Pantaleon *et al*., 2018, Poquet *et al*. 2018). Dapa *et al*. showed that strains 630 (ribotype 012) and R20291 (ribotype 027) could form biofilms after 1 and 3 days of growth in rich BHIS glucose medium in microplates. Dawson *et al*. (2021) confirmed this observation for strains 630 and R20291 and showed that a representative strain of three other ribotypes, strains CD305 (ribotype 023), M68 (ribotype 017) and M120 (ribotype 078), were also able to form biofilms under the same conditions. In *Bacillus subtilis*, non-domesticated compared to model strains were shown to display original biofilm properties (Bridier *et al*., 2011). The study of *C. difficile* non-domesticated (or ’wild-type’) strains for their capacity to form biofilms is more recent. In 2018, Pantaléon et al. analysed thirty-seven laboratory strains of nineteen different ribotypes, including RT126, RT078, RT020 and RT005. All strains were capable of forming biofilms of varying density and adhesiveness (Pantaléon et *al*., 2018). Morais *et al* studied three strains of ribotype 027 isolated from patients in South America. They were able to form biofilms, which, after 48 hours of growth, were of similar biomass (Morais *et al*., 2022). Similarly, Wultanska *et al* observed that twelve clinical strains from four different ribotypes (RT017, RT023, RT027 and RT176) formed biofilms after 48 hours. Their study also revealed different biofilm densities, with RT027 strains forming denser biofilms compared to those formed by strains of ribotypes RT017, RT023 and RT176 (Wultanska *et al*., 2023).

We analysed the biovolume of intact biofilms, which were observed *in situ* after live/dead staining by CLSM and the adhesive and wash-resistant biomass of biofilms stained by CV. This showed that the non-toxigenic strains of ribotype WRT009 formed more adhesive and resistant to wash biofilms, i. e. more cohesive biofilms. These strains could be promising in the future. Indeed, several non-toxigenic strains, including some of ribotype 009, have already been shown to coexist with toxigenic strains in different animal hosts, such as rodents, piglets, companion animals and horses (Gerding *et al*., 2015; Arruda *et al*., 2016; Gerding *et al*., 2018). Moreover, non-toxigenic strains have been shown to display a protective effect against toxigenic strains. Merrigan et al. used a hamster model to study CDI prevention by non-toxigenic strains. Hamsters continuously treated with clindamycin were given one non-toxigenic strain, either a clindamycin-resistant (M13) or a susceptible one (M3), and challenged by a toxigenic strain. All hamsters were colonised by the resistant non-toxigenic strain and protected from CDI by the toxigenic strain. Moreover, when the susceptible non-toxigenic strain was given after clindamycin withdraw, it was also able to colonise and protect hamsters from the challenging toxigenic strain (Merrigan *et al*., 2003). More recently, non-toxigenic strains of ribotypes RT032, RT010, RT140, RT151, RT388, RT031 and RT847 collected from premature infants have also been shown to prevent colonisation by toxigenic strains (RT027) and reduce mortality in hamsters (Couturier *et al*., 2020). Importantly, a non-toxigenic RT009 strain not only prevented CDI in model piglets experimentally challenged by a toxigenic strain (Oliveira Junior *et al*., 2019a), but also reduced the incidence of CDI in piglets from a commercial farm (Oliveira Junior *et al*., 2019b). In this context, as biofilms formed by non-toxigenic strains of ribotype 009, compared to those formed by other strains, were shown here to be more cohesive and adhesive to the abiotic device, it would be interesting to evaluate whether this property is correlated to a better colonisation of the gut *in vivo*. Of note, strain *Cd* E-61-CDL of WRT009 was among the strains whose biofilm viability, in contrast to planktonic viability, was unaffected even after an early addition of vancomycin, suggesting that this increased tolerance could contribute to persistence and therefore to a very early effect of protection against a contaminating toxigenic strain.

Our microscopic analyses, including xy images, showed that all biofilms had a dense structure. A similar structure has previously been observed for biofilms of laboratory strains 630 and R20291 after grown for 48 h in BHIS medium in microplates (Dapa *et al*., 2013), and those of strains BI17 and BY1 grown for 48 h in tryptic soy medium on polycarbonate membranes (Semenyuk et al., 2014). We observed that the biofilm structure of our non-domesticated strains was very similar to that of our laboratory reference, 630Δ*erm*, as well as to that of its parental strain, 630 cultured in a rich BHIS glucose medium for 24 to 72h (Dapa *et al*., 2013).

A study of *Bacillus subtilis* revealed notable differences between laboratory strains and non-domesticated strains isolated from food or medical environments. Non-domesticated strains produced larger and more irregular biofilms, whereas laboratory strains formed smaller and more homogeneous biofilms (Bridier *et al*., 2011). Undomesticated strains of *C. difficile* might differ from those of *B. subtilis* according to the ability to form biofilms of very diverse architectures. Alternatively, the conditions for biofilm formation, in a rich medium supplemented by glucose, may not have allowed revealing significant differences. We previously observed a very different biofilm architecture, high, sparse and heterogeneously made of individual and micro-aggregated cells, after growth of the laboratory strain 630Δ*erm* in TYt medium. It could therefore be interesting to test whether undomesticated strains could form biofilms of different architectures in TYt (Poquet et *al*., 2018).

We examined *in vitro* biofilms formed by undomesticated strains for properties that could be pertinent *in vivo,* during the infection cycle. Production of the toxins TcdA and TcdB is essential for virulence, and spore production is important for dissemination of *C. difficile* (Frost *et al*., 2021). Under our experimental conditions, after 48 hours of growth in a BHIS glucose medium, we did not detect any production of TcdA or TcdB, neither in biofilms nor in planktonic cultures. In a previous study using clinical *C. difficile* strains K14, BI6 (RT027), VPI10463 (RT087), J9, 630 (RT012), BI17 (RT027) and BY1, toxins could be detected by Elisa in biofilms grown in tryptic soy medium for three days (Semenyuk et *al*., 2014). However, other studies have also shown that toxin gene expression was repressed in biofilms compared to planktonic cultures. In strain R20291 grown for 7 days in BHIS medium either as a biofilm on glass beads or as a planktonic culture, an RNA-seq analysis showed that *tcdB* gene was repressed (by 2.8-fold) in biofilms compared to planktonic cultures in the same condition (Maldarelli *et al*., 2016). In 630Δ*erm* grown in TY medium in a continuous flow biofilm compared to a planktonic culture, *tcdA* expression was found to decrease (by 25-fold), while no significant change in *tcdB* expression was detected (Poquet *et al*., 2018). The differences in growth conditions and strains make direct comparisons between studies difficult.

We observed a low sporulation rate ranging from 0.5% to 10% in both biofilms and planktonic cultures. In strains 630, VPI, K14, BI17, BI6, J9 and BY1, biofilms after six days of growth in TY medium on polycarbonate membranes showed sporulation rates ranging from about 9% to 70% depending on the strains (Semenyuk *et al*., 2014). Pickering *et al*. showed significant differences in spore formation within biofilms aged 7 to 10 days formed in BHI medium tubes for strains of ribotypes RT001, RT020, RT027 and RT078 (Pickering *et al*., 2018). Dawson et *al*. also observed that spore production by strains 630E and R20291 was very low in 3-day-old biofilms (∼ 10^3^ spores/mL), and increased in 6-day-old biofilms, reaching around 10^6^ spores per millilitre (Dawson et *al*., 2012). Therefore, sporulation seemed to be a property of old biofilms. In our study, low sporulation is therefore not unexpected in two days-old biofilms grown in rich medium in the presence of glucose. Plaza-Garrido et al. (2015) studied two laboratory strains (630 and R20291) and seventeen non-domesticated strains isolated from patients with recurrent or non-recurrent CDI, and their biofilms after five days of growth in BHIS medium in the presence or not of added glucose at the same concentration as we used. They showed sporulation percentages ranging from 0.0001% to 100%, depending on whether the strains were laboratory or non-domesticated, with laboratory strains generally showing high sporulation rates. They also found that glucose significantly reduced sporulation in clinical isolates during biofilm development (Plaza-Garrido *et al*., 2015). Furthermore, a recent study by Wetzel *et al*. demonstrated the effect of pH in BHIS taurocholate medium on the sporulation capacity of planktonic cultures. Under acidic conditions, strains 630Δ*erm* and R20291 produced significantly fewer spores and even no spores at pH=5.2 (Wetzel et McBride., 2020). In our experiments, we measured the pH of the biofilm supernatants after 48 hours of growth in BHIS glucose medium. The pH of the supernatants was acidic, ranging from 5 to 6.5, in agreement with the low sporulation rate we observed.

We then investigated the tolerance of biofilms and planktonic cultures to vancomycin, which is recommended as a second-line treatment for CDI (van Prehn *et al*., 2021). We added vancomycin at the same concentration to the biofilm and the planktonic culture of each strain and compared cell viability after vancomycin exposure or not. We used a concentration of 8 MIC for each strain, i. e. a concentration ranging from 1 to 8mg/L (Table 1), and of the same order of magnitude as the estimated concentration resulting from the maximal vancomycin concentration in the colon (assuming its volume to be of ∼2L), after treatment by one oral dose of 125 mg (which has to be added four times a day to a patient) (van Prehn *et al*., 2021).

Our study showed that early administration of vancomycin at 8 MIC significantly reduced the viability of all nine studied strains (eight non domesticated ones and the reference) in planktonic cultures, but of only five of them in biofilms, and the reduction level was much higher in the planktonic culture than in the biofilm of the same strain. Late administration had no effect. Previous studies investigated the tolerance of biofilms to antibiotics (Dawson *et al*., 2021; Morais *et al*., 2022). In the first study, the planktonic phase was removed from three-day-old biofilms grown in BHIS glucose medium in microplates, and the remaining (adhesive) biofilm was allowed to grow for a further 48 hours after being resuspended in medium containing vancomycin at different inhibitory concentrations (1xMIC, 4xMIC and 8xMIC). This study showed a vancomycin dose-dependent decrease in the biomass of biofilms stained by crystal violet (Morais et *al*., 2022). In Dapa et *al* 2013, the significant resistance of *C. difficile* biofilms was demonstrated using a 100x MIC of vancomycin, showing that biofilm bacteria survived 5 and 12 times more than planktonic bacteria, depending of growth length, 1 day and 3 days respectively. This suggested that biofilms could play a protective role against antibiotic treatment (Dapa et *al*., 2013). In Dawson *et al*., the 3- day-old biofilm of strain 630 grown in BHIS medium was examined after vancomycin addition at a concentration of 12.5 mg/mL, i. e. at least 3 orders of magnitude higher than either the concentrations we used or the MIC of resistant *C. difficile* strains (≥4mg/L, Krutova et *al*., 2022). In biofilms, after addition to such a very high concentration, the viable vegetative cell count was only reduced by a ten-fold factor, reaching 7.7% of the untreated level (Dawson *et al*., 2021). Vancomycin effect was limited, demonstrating the ability of *in vitro* biofilms to tolerate huge concentrations of antibiotics. Biofilm tolerance to antibiotics is very well documented in many other pathogens (Brooun et *al*., 2000; Spoering et Lewis., 2001; Gu et *al*., 2019; Kamble et Pardesi., 2020; Yonezawa et *al*., 2013). The higher tolerance of biofilms than of planktonic cultures (Brooun et *al*., 2000, Spoering et Lewis., 2001) could be attributed to several factors, such as the presence of extracellular DNA (eDNA) or extracellular polymeric substances (EPS), which can form a physical and chemical barrier preventing effective penetration of antibiotics or the existence of phenotypically tolerant or persister cells (Rubio-Mendoza *et al*., 2023).

Our study showed that *C. difficile in vitro* biofilms were highly tolerant to a realistic, therapeutic dose of vancomycin. If the biofilms studied here were relevant *in vivo*, they might contribute to the difficulty to completely eradicate *C. difficile* and finally the possibility of re-infection after the end of treatment.

In conclusion, our study showed that non-domesticated *C. difficile* strains isolated from equids formed highly similar biofilms after 48h of growth in BHIS glucose medium than the model strain. Of note, the biofilms of non-toxigenic strains of ribotype WRT009 were of smaller biovolume, but higher adherence. Finally, vancomycin at a realistic concentration was more efficient when added at an early rather than late growth stage, and much more efficient against planktonic cultures than biofilms.

Supplementary Data

## Acknowledgements

Authors thank the Mima2 platform (INRAe, Institut Micalis) for their imaging facilities and their training in the use of the confocal microscope. The authors also thank Lucas Bertrand for writing the computer scripts for the CLSM analyses. We would like to acknowledge Olga Soutourina, Claire Janoir, Christina Nielsen Leroux, Imad Kansau, Romain Briandet and Julien Deschamps for helpful discussions. We are deeply grateful to Christina Nielsen Leroux for her constant support during the during course of this work.

## Author contributions

A.L. and I.P. designed the research. A.L. performed the experiments. LB helped in several technical aspects, including image analysis (script for Image J). A.L. and I.P. analysed the data. A.L. and I.P. wrote and edited the manuscript. S.P. and I.P. provided the collection of unpublished, deeply characterized C. difficile strains originating from necropsied equids. V.C. trained AL in CLSM on MIMA2 platform of INRAE. A.C. trained and helped AL in ELISA tests. IP wrote and submitted a PhD project to Ecole Doctorale Abies from University Paris-Saclay, and Abies selected it for the PhD grant competition from the French government (Ministère Supérieur de l’Enseignement et de la Recherche) in 2022. All authors read and agree to the submitted version of the manuscript.

