## Supplementary data for "*In vitro* biofilms of *Clostridioides difficile* undomesticated strains: Morphology and properties according to strain diversity"

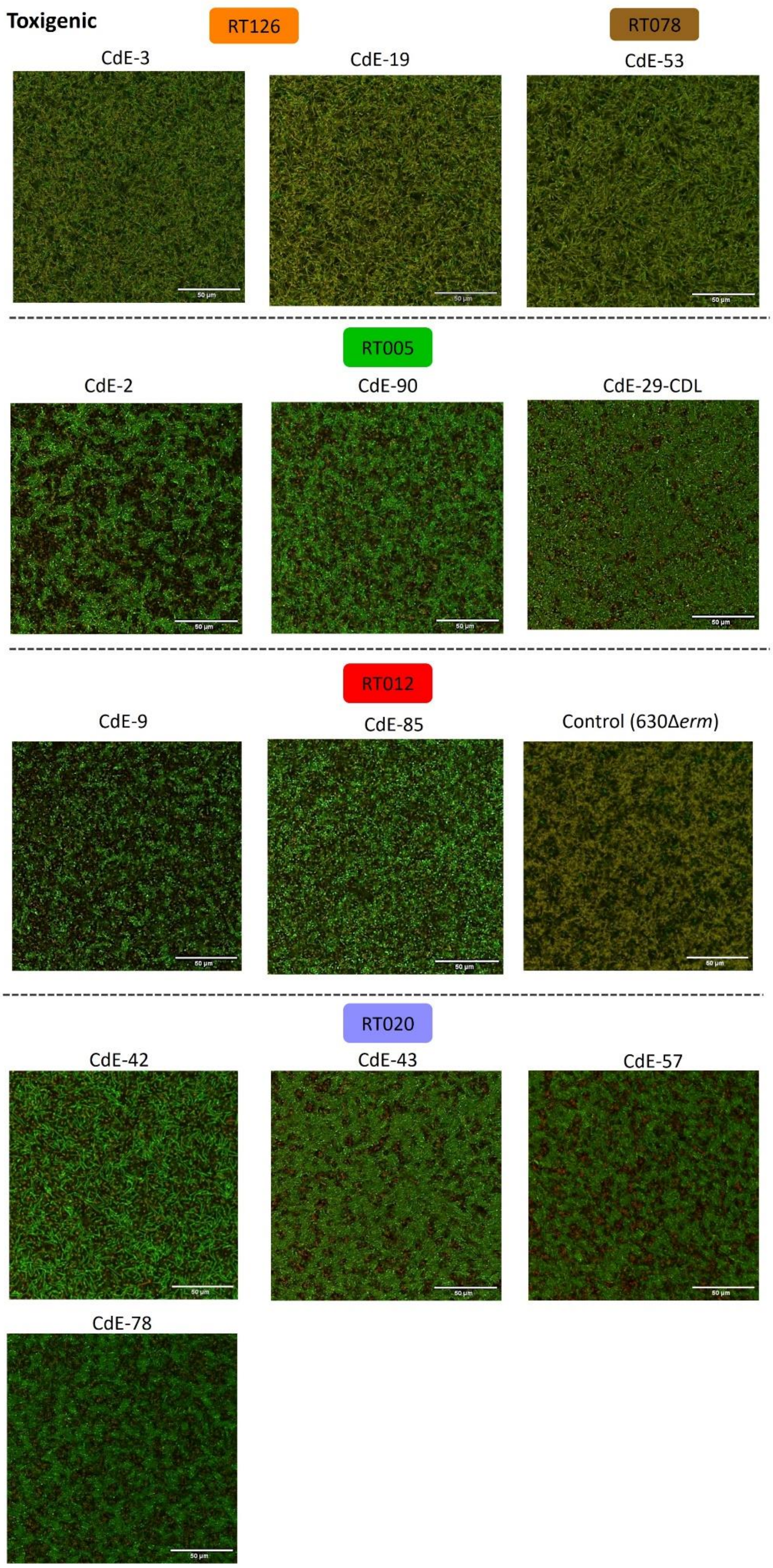
Toxigenic
RT126
RT078
CdE-3
CdE-19
CdE-53
RT005
CdE-2
CdE-90
CdE-29-CDL
RT012
CdE-9
CdE-85
Control (630Δ*erm*)
RT020
CdE-42
CdE-43
CdE-57
CdE-78

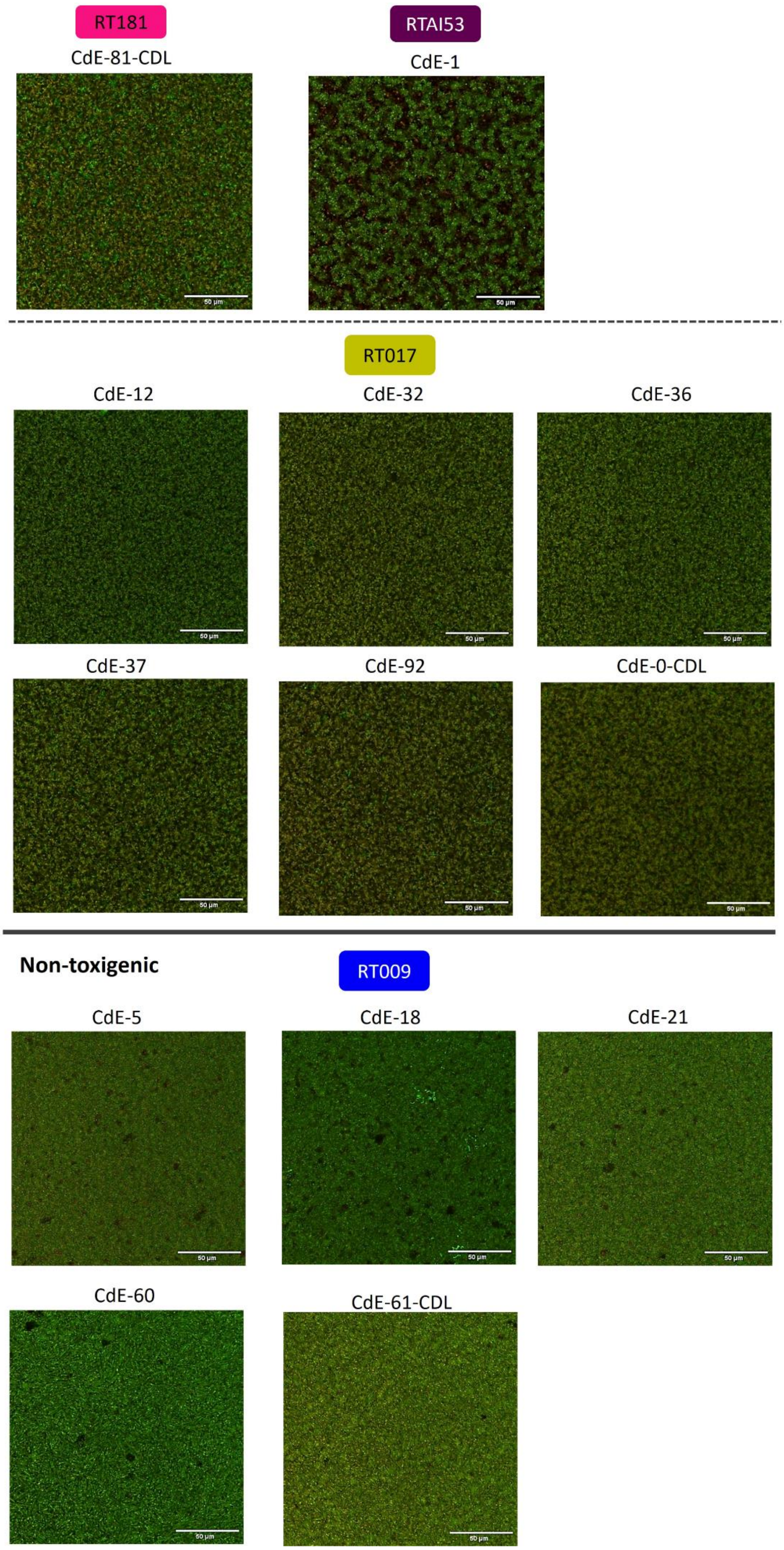
RT181
CdE-81-CDL
RTAI53
CdE-1
RT017
CdE-12
CdE-32
CdE-36
CdE-37
CdE-92
CdE-0-CDL
Non-toxigenic
RT009
CdE-5
CdE-18
CdE-21
CdE-60
CdE-61-CDL

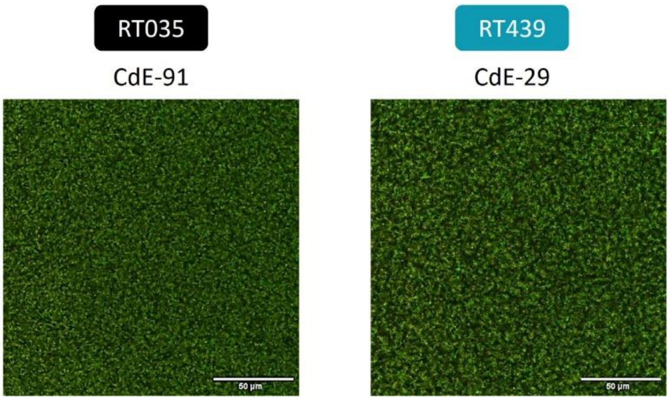


**Supplementary data 1** : Biofilm structures in xy image. Twenty-seven equine *C. difficile* strains and the reference strain 630Δ*erm* were studied. Biofilms were observed after live-dead staining by CLSM. xy images at z= 25 were extracted from biofilm stacks using ImageJ. Observations were repeated 3 times for each equine strain.

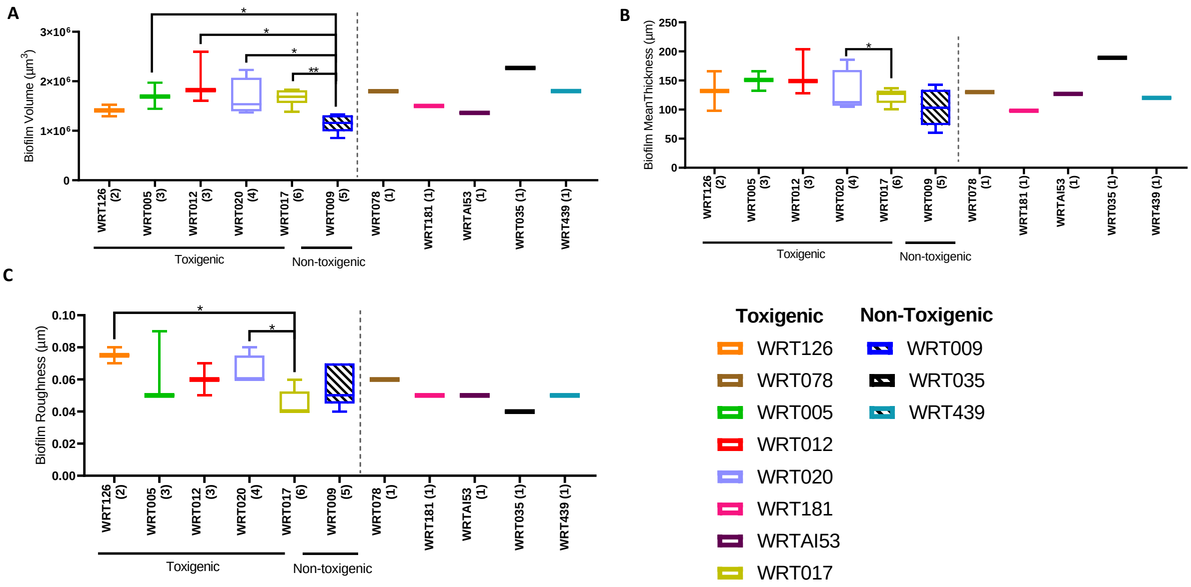


**Supplementary data 2**: Biofilm 3D structure analysed according to strain PCR-ribotype. After live dead staining and observation of biofilms by CLSM, their structure was evaluated using BiofilmQ software. The experiment was repeated 3 times for each equine strain. The results were analysed according to strain ribotype. (A) Biofilm biovolume (total fluorescence). (B) Biofilm mean thickness. (C) Biofilm roughness is defined as the standard deviation of thickness across different vertical sections of the biofilms. Wilcoxon-Mann-Whitney test * p<0.05, ** p<0.01, *** p<0.001.

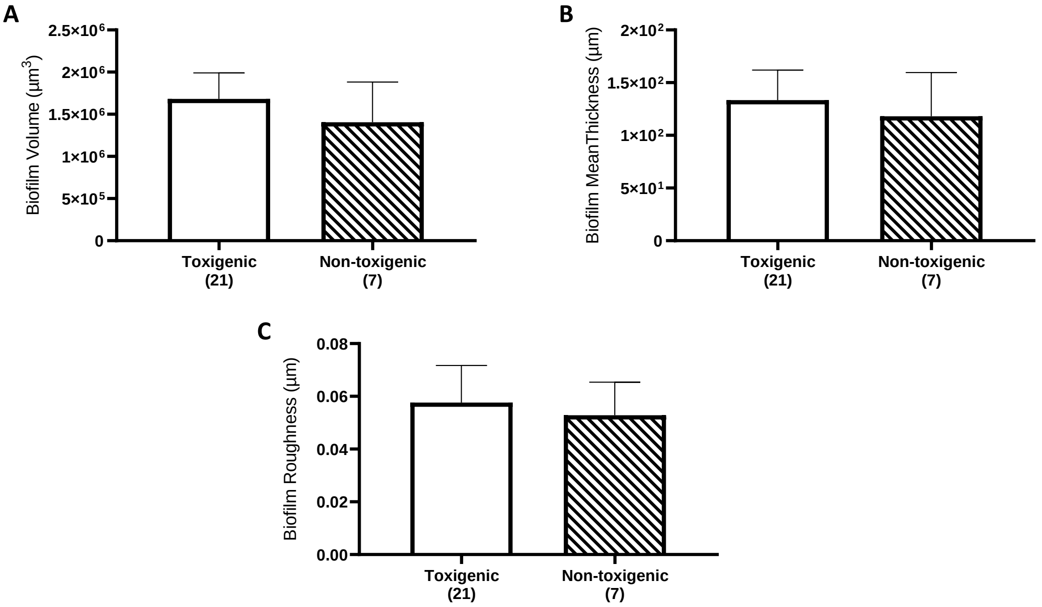


**Supplementary data 3**: Biofilm structure based on pathogenicity. We studied 27 equine strains of *C. difficile* together with the laboratory strain 630Δ*erm* used as a reference. Biofilms were stained with Syto9 (for live bacteria) and propidium iodide (for damaged and dead bacteria). They were observed by CSLM and their structure was evaluated using BiofilmQ software. The experiment was repeated 3 times for each equine strain. The strains were grouped according to their pathogenicity. (A) Biovolume. (B) Mean thickness. (C) Roughness. (D) Percentage of cell death. Wilcoxon-Mann-Whitney test: not significant.

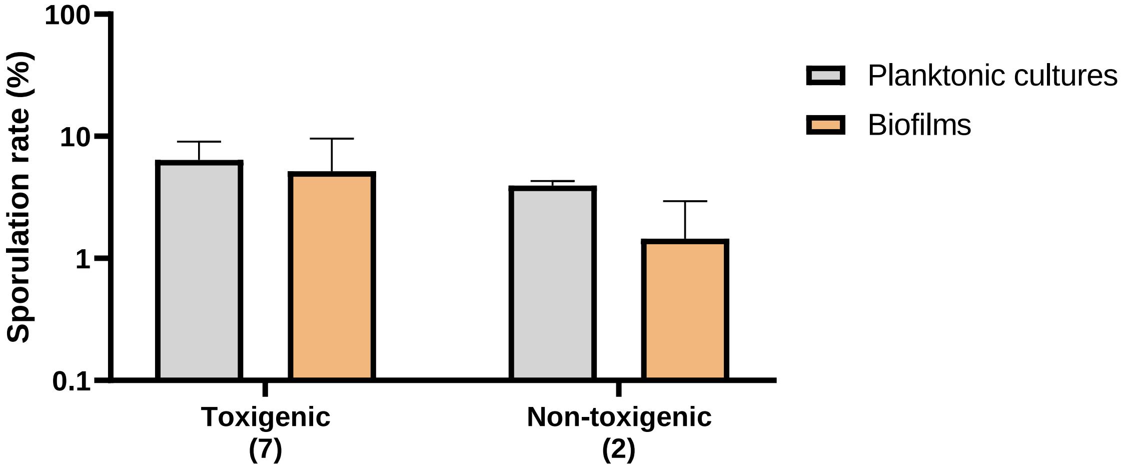


**Supplementary data 4**: Sporulation rate based on pathogenicity of planktonic cultures and biofilms. We studied 8 equine strains of *C. difficile* together with the reference laboratory strain 630Δ*erm*. The strains were grouped according to their pathogenicity. The percentage of spores was calculated as the number of heat-resistant spores relative to the total cell count. The results are presented on a semi-logarithmic scale. The experiment was repeated four times for each strain. Wilcoxon-Mann-Whitney test: not significant.

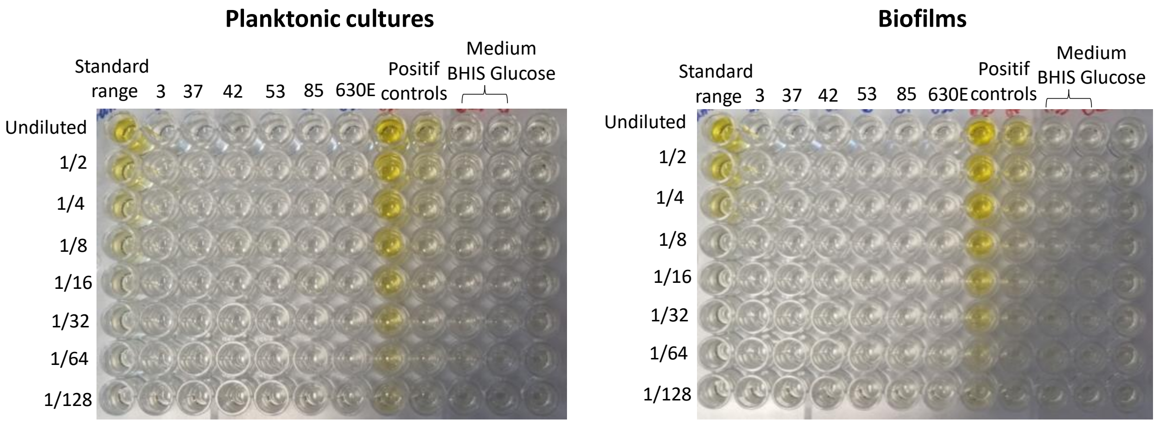


**Supplementary data 5**: TcdB production of planktonic cultures and biofilms. TcdB production quantified by ELISA for the 5 toxigenic strains and the reference laboratory strain 630Δ*erm*. The experiment was repeated 2 times for each strain.
